# Slow Wave Sleep Reduces CSF Concentrations of Beta-amyloid and Tau: A Randomized Crossover Study in Healthy Adults

**DOI:** 10.1101/2025.06.15.659740

**Authors:** Tim Lyckenvik, Martin Olsson, My Forsberg, Pontus Wasling, Henrik Zetterberg, Jan Hedner, Eric Hanse

**Affiliations:** Department of Physiology, Institute of Neuroscience and Physiology, Sahlgrenska Academy at the University of Gothenburg, Gothenburg, Sweden; Department of Neurology, Sahlgrenska University Hospital, Gothenburg, Sweden; Department of Internal Medicine and Clinical Nutrition, Institute of Medicine, Sahlgrenska Academy at the University of Gothenburg, Gothenburg, Sweden; Department of Anesthesiology and Intensive Care, Sahlgrenska University Hospital/Östra, Gothenburg, Sweden; Department of Clinical Neuroscience, Institute of Neuroscience and Physiology, Sahlgrenska Academy at the University of Gothenburg, Gothenburg, Sweden; Department of Psychiatry and Neurochemistry, Institute of Neuroscience and Physiology, Sahlgrenska Academy at the University of Gothenburg, Mölndal, Sweden; Wisconsin Alzheimer’s Disease Research Center, School of Medicine and Public Health, University of Wisconsin-Madison, Madison, United States; Clinical Neurochemistry Laboratory, Sahlgrenska University Hospital, Mölndal, Sweden; Department of Neurodegenerative Disease, UCL Institute of Neurology, London, United Kingdom; UK Dementia Research Institute at UCL, London, United Kingdom; Hong Kong Center for Neurodegenerative Diseases, Clear Water Bay, Hong Kong, China; Centre for Brain Research, Indian Institute of Science, Bangalore, India

**Keywords:** Alzheimer’s disease, sleep, slow wave sleep, tau, amyloid beta

## Abstract

**Background:** Slow-wave sleep has been proposed to facilitate the removal of proteins, implicated in neurodegeneration, from the brain. While mechanistic evidence from animal models is accumulating, direct human data on how slow-wave sleep shapes cerebrospinal fluid (CSF) proteostasis remain limited, constraining our understanding of physiological resilience to neurodegenerative disease.

**Methods:** Twelve healthy adults (aged 20–40 years) underwent CSF sampling following three controlled sleep conditions in a randomized crossover design; (1) one night of sleep followed by afternoon CSF sampling, (2) one night of sleep followed by morning CSF sampling, and (3) one night of total sleep deprivation followed by morning CSF sampling.

Sleep and wakefulness were verified using polysomnography and actigraphy, with >4-week washout periods between conditions.

Measured CSF biomarkers included Alzheimer’s disease-related proteins: beta-amyloid isoforms (Aβ38, Aβ40, and Aβ42), total and phosphorylated tau, glial fibrillary acidic protein (GFAP), and neurofilament light, as well as orexin, albumin (also measured in serum), and osmolality. Differences between conditions were assessed using Friedman tests with Dunn’s post hoc correction.

**Results:** CSF levels of Aβ and tau tended to be consistently lower after sleep compared with both afternoon sampling and post-sleep deprivation. Concurrently, CSF albumin levels increased after sleep, while neurofilament light and GFAP remained unchanged. Orexin levels rose markedly during sleep deprivation but showed no circadian variation.

**Conclusions:** These findings support a model in which slow wave sleep enhances CSF turnover, reducing concentrations of specific proteins, including Aβ and tau. Understanding how sleep regulates the homeostasis of neurodegeneration-related proteins may inform strategies to mitigate disease progression.

## Background

Concentration-dependent aggregation of neurotoxic variants of endogenous proteins is implicated in several neurodegenerative diseases, including Alzheimer’s disease (AD) and Parkinson’s disease (PD) [1]. Two of the key proteins implicated in pathological aggregation in AD, beta-amyloid (Aβ) [2, 3] and tau [4-6], are physiologically released during synaptic activity, making efficient clearance potentially essential to prevent neurotoxic accumulation.

The pathological hallmark of AD is the accumulation of misfolded Aβ and tau proteins, which form senile plaques and neurofibrillary tangles, respectively [7]. Senile plaques consist primarily of the aggregation-prone Aβ42 isoform, whereas tangles are composed of hyperphosphorylated tau. Experimental studies in healthy individuals have shown that total sleep deprivation (TSD) leads to elevated morning concentrations of Aβ42 [8-10] and tau [11, 12] in cerebrospinal fluid (CSF), supporting a role of sleep in proteostatic balance. In addition, disrupted sleep patterns have been associated with increased AD risk [13, 14], suggesting that impaired clearance during sleep may contribute to the accumulation of Aβ42 and tau over time.

According to the glymphatic hypothesis on brain metabolite, peptide and protein clearance [15], sleep and general anesthesia facilitate convective fluid movement in the brain parenchyma, enhancing perivenous clearance of neuroactive substances such as Aβ [16], although this concept is actively debated [17-19]. In a previous study, we reported that restricting sleep to less than four hours per night for five consecutive nights did not significantly affect CSF biomarkers for amyloid accumulation or AD-type neurodegeneration, even though the time spent in all sleep stages except SWS was reduced [20]. These findings suggest that the preservation of slow-wave sleep (SWS) may be sufficient to maintain short-term proteostatic balance despite overall sleep loss.

Moreover, parenchymal clearance is increased by multisensory-evoked synchronous oscillation of neuronal activity at low frequencies, such as those observed during SWS [21]. Increased clearance has also been observed at higher frequencies [22] not associated with sleep, suggesting that the beneficial features of SWS may be therapeutically separable from sleep *per se*.

We hypothesized that eliminating SWS by total sleep deprivation impedes the physiological clearance of CSF biomarkers associated with AD, in contrast to the effects of partial sleep deprivation. To test this hypothesis, we examined the impact of one night of TSD on CSF concentrations of biomarkers reflecting key aspects of AD pathophysiology by measuring the Aβ isoforms Aβ38, Aβ40 and Aβ42 and hyperphosphorylated tau (p-tau) together with total tau (t-tau). Neuronal injury was further assessed by neurofilament light chain (NfL), whereas astroglial activation was gauged by glial fibrillary acidic protein (GFAP). Finally, the CSF orexin concentration was measured because of its critical role in sleep regulation and its proposed association with AD pathology [27].

By exploring these biomarkers, our study aims to clarify how SWS affects the clearance of proteins implicated in AD to potentially uncover targetable mechanisms for mitigating neurodegeneration.

## Methods

### Design

This study uses the samples collected in Forsberg et al. (2021) [28], which was a randomized, crossover study in which each participant was exposed to three different sleep conditions at the sleep laboratory, separated by at least four weeks, to allow within-subject comparisons of CSF biomarker levels after sleep and sleep deprivation (Figure 1). Blood and CSF were sampled after each night, either in the morning at 6-7 a.m. following sleep or sleep deprivation, or in the afternoon at 3-5 p.m. following nighttime sleep. The order of the sampling conditions was randomized, and an experienced neurologist collected 10 ml of lumbar CSF on each occasion. Actigraphy was used to ensure that wakefulness had been sustained in the sleep deprivation group, while sleep was monitored with polysomnography (PSG) the night before sampling for both other groups, as previously described [28].

**Figure 1.**
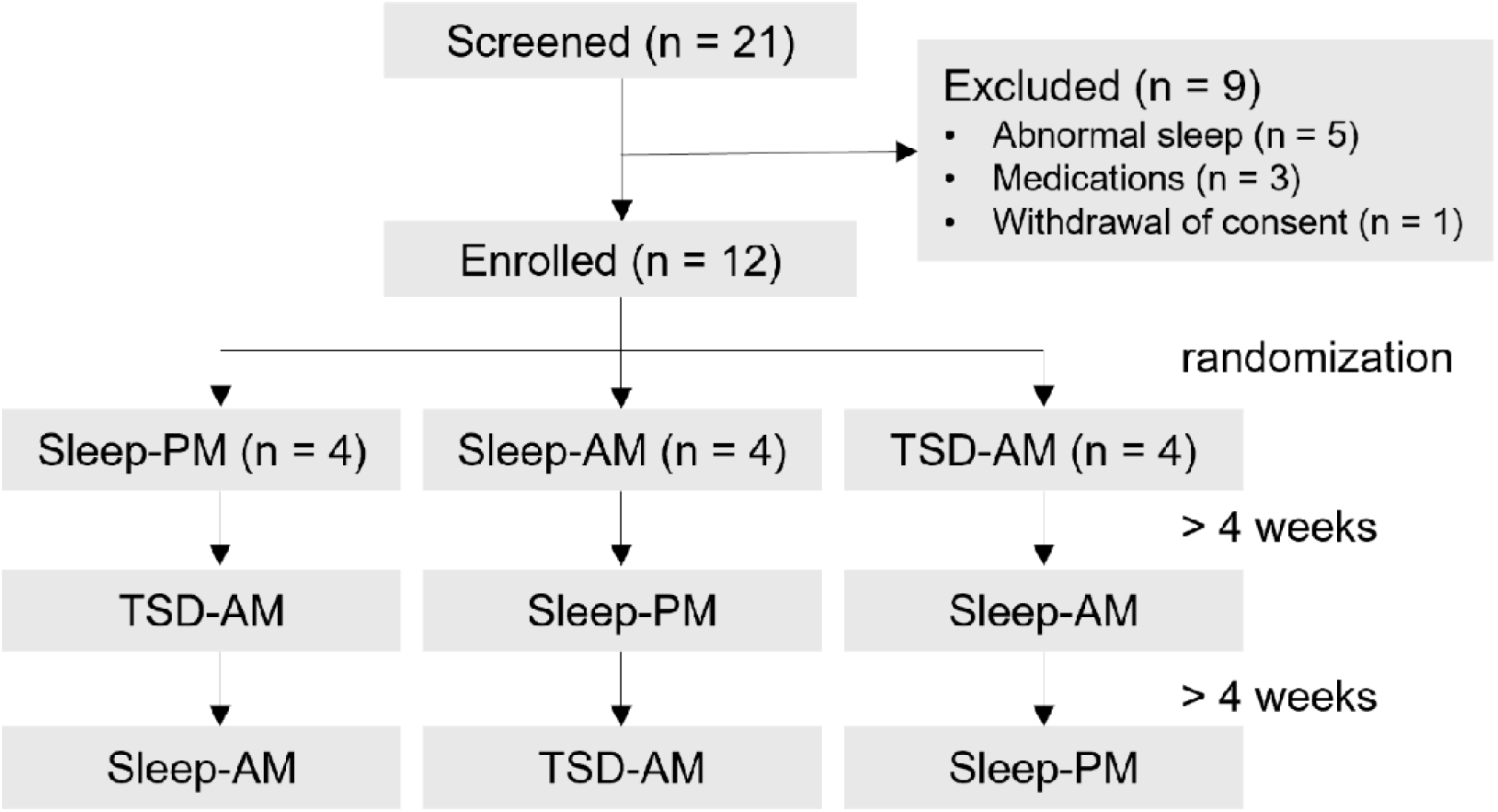
Study flow and condition order according to a randomized crossover design. Enrolled participants (n=12) were randomized into three sequences of sleep conditions. Each condition was separated by ≥4 weeks. Abbreviations: Sleep-PM, afternoon samples after nighttime sleep; Sleep-AM, morning samples after sleep; TSD-AM, morning samples after total sleep deprivation.

Participants were instructed to refrain from excessive sleep in preparation for nights in the sleep laboratory and to avoid the intake of drugs (such as caffeine and nicotine) or alcohol that may affect sleep) for at least 24 hours before the experiments. Furthermore, the participants carried actigraphy recorders for 24 hours prior to arriving in the sleep laboratory on each occasion as well as during the entire sleep-deprivation night to ensure the exclusion of participants with sustained sleep periods preceding sleep-deprivation sampling. Prior to this sampling, the participants spent the night in the laboratory with lights on playing games, studying, talking, and walking around, refraining from more strenuous physical activity.

Morning CSF was collected following sleep from participants who remained in bed with the lights out in individual rooms for 8 hours until the completion of sampling. During the day preceding the afternoon sampling, participants sustained unmonitored wakefulness outside the laboratory. There was a loss of PSG data preceding one of the 12 afternoon samplings.

### Participants

Twenty-one volunteers were screened for general health and sleep habits, of which 12 healthy individuals (seven females, five males) were included. Nine subjects were excluded during screening because of abnormal sleep (5), medications (3) or withdrawal of consent (1). The inclusion criteria were as follows: habitual sleep duration of 6–9 hours in the time window between 9 p.m. and 8 a.m. with sleep latency < 30 min. The exclusion criteria were as follows: nocturnal awakenings, restless legs syndrome, excessive daytime sleepiness (evaluated with the Epworth Sleepiness Scale > 10), morning headache and dryness of the mouth [29, 30]. Participants were also excluded if they had worked a nightshift or traveled across more than two time zones within 2 months of CSF collection.

### Biomarkers

All measurements were conducted by board-certified laboratory technicians at the Clinical Chemistry Laboratory at Sahlgrenska University Hospital. The laboratory is accredited under the Swedish Accreditation body (SWEDAC).

Amyloid β (Aβ) proteins (Aβ38, Aβ40 and Aβ42) were measured using electrochemiluminescence assays (Meso Scale Discovery, Rockville, Maryland, United States). Total tau (t-tau) and phosphorylated tau (p-tau) were measured using Lumipulse technology (Fujirebio, Ghent, Belgium). Neurofilament light chain (NfL) and GFAP were measured using validated in-house ELISA methods [31-33]. Orexin-A levels in CSF were measured using an in-house radioimmunoassay (RIA), with a normal reference range defined as >400 pg/mL [34]. CSF and serum albumin concentrations were measured via immunonephelometry using a Beckman Image Immunochemistry system (Beckman Instruments, Beckman Coulter, Brea, CA, USA). Qalb was calculated as CSF albumin (mg/L) divided by serum albumin (g/L). Osmolality was measured as previously described [35]. All protein concentration measurements were performed in one round of experiments using one batch of reagents with intra-assay coefficients of variation <10%.

As previously described [28], the concentrations of Na^+^, K^+^ and Cl^-^ were measured using ion-selective electrodes (ISEs), whereas the Ca^2+^ and Mg^2+^ concentrations were determined colorimetrically using the o-cresolphthaleion and chlorophonazo III methods, respectively.

### Statistical analysis

The data were analysed, and graphs were created using the software GraphPad Prism^®^ (GraphPad Software, version 10.4.1 (627)). The statistical significance of differences between groups was determined by paired one-way ANOVA (Friedman test) with Dunn’s test for correction for multiple comparisons. Spearman correlation analyses with associated p values were used to visualize correlations between the biomarkers under each condition. Data are presented as the median (IQR).

## Results

The results from analyses of polysomnography, CSF ion concentrations, and albumin concentrations and osmolality in both serum and CSF have been published previously [28].

### Decreased levels of Aβ and tau, but not NfL or GFAP, after one night of sleep

The concentrations of tau, particularly p-tau, and the measured isoforms of Aβ (Aβ38, Aβ40 and Aβ42) followed a consistent trend of being lowest in the samples collected in the morning following a full night of sleep (Figure 2A-C, E-F). NfL and GFAP remained stable (Figure 2G-H). Aβ40 and Aβ42 were found in similar proportions across conditions, as reflected by the stability of their ratio (Figure 2D).

**Figure 2.**
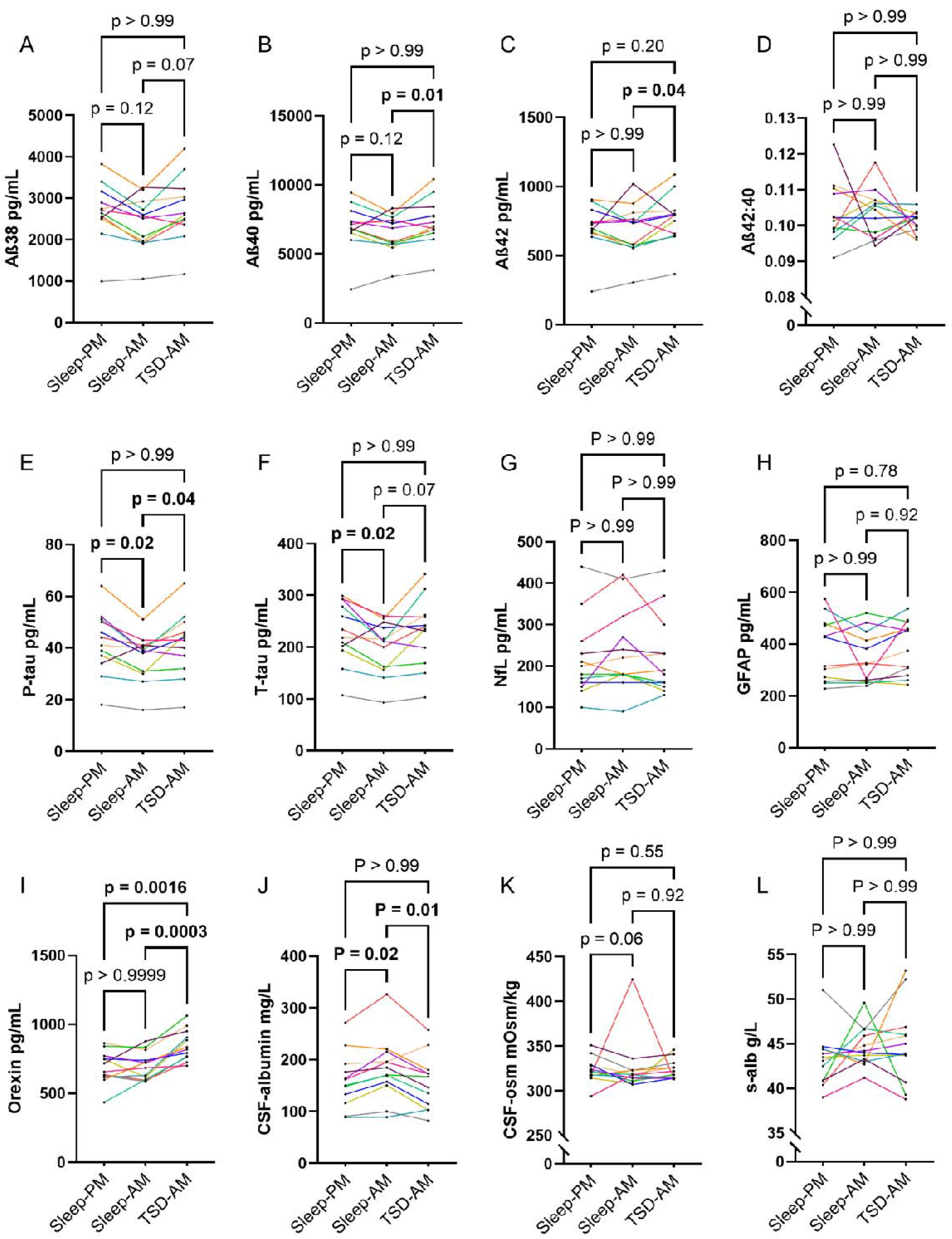
Decreased levels of Aβ and tau, but not NfL or GFAP, after one night of sleep. Biomarker concentrations across conditions. Each line represents paired measurements from a single individual across the three conditions. Abbreviations: Sleep-PM, afternoon samples after nighttime sleep; Sleep-AM, morning samples after sleep; TSD-AM, morning samples after total sleep deprivation; Aβ38, amyloid β (1-38); Aβ40, amyloid β (1-40); Aβ42, amyloid β (1-42); Aβ42:40, ratio of beta amyloid 42/40; T-tau, total tau; P-tau, phosphorylated tau; NfL, neurofilament light chain; GFAP, glial fibrillary acidic protein; CSF-osm, CSF osmolality; s-alb, serum albumin concentration. The median values and interquartile ranges are provided in Supplementary Table 1.

Aβ40, Aβ42 and p-tau concentrations were significantly lower in the morning samples collected after sleep than in the samples collected after total sleep deprivation. Differences in Aβ38 and t-tau did not reach statistical significance. Similarly, p-tau and t-tau concentrations were significantly lower in samples collected in the morning after sleep than in those collected in the afternoon, whereas the corresponding differences in Aβ were not statistically significant because samples from two individuals exhibited the opposite pattern (Figure 2A-C).

Orexin levels were significantly elevated in morning samples following sleep deprivation compared with both morning samples after sleep and afternoon samples (Figure 2I).

### Increased CSF albumin after one night of sleep

CSF albumin showed an inverse pattern to Aβ and tau, with higher concentrations in the morning after sleep compared with both sleep deprivation and afternoon sampling (Figure 2J). This difference was not explained by serum albumin, which remained stable across conditions (Figure 2L). Similarly, CSF osmolality showed a diurnal pattern that did not differ significantly between conditions, possibly due to a high outlier measured in the morning after sleep (Figure 2K). However, osmolality was not consistently correlated with albumin or any other measured biomarkers across conditions (Figure 3), suggesting that osmolality did not contribute meaningfully to the biomarker differences.

**Figure 3.**
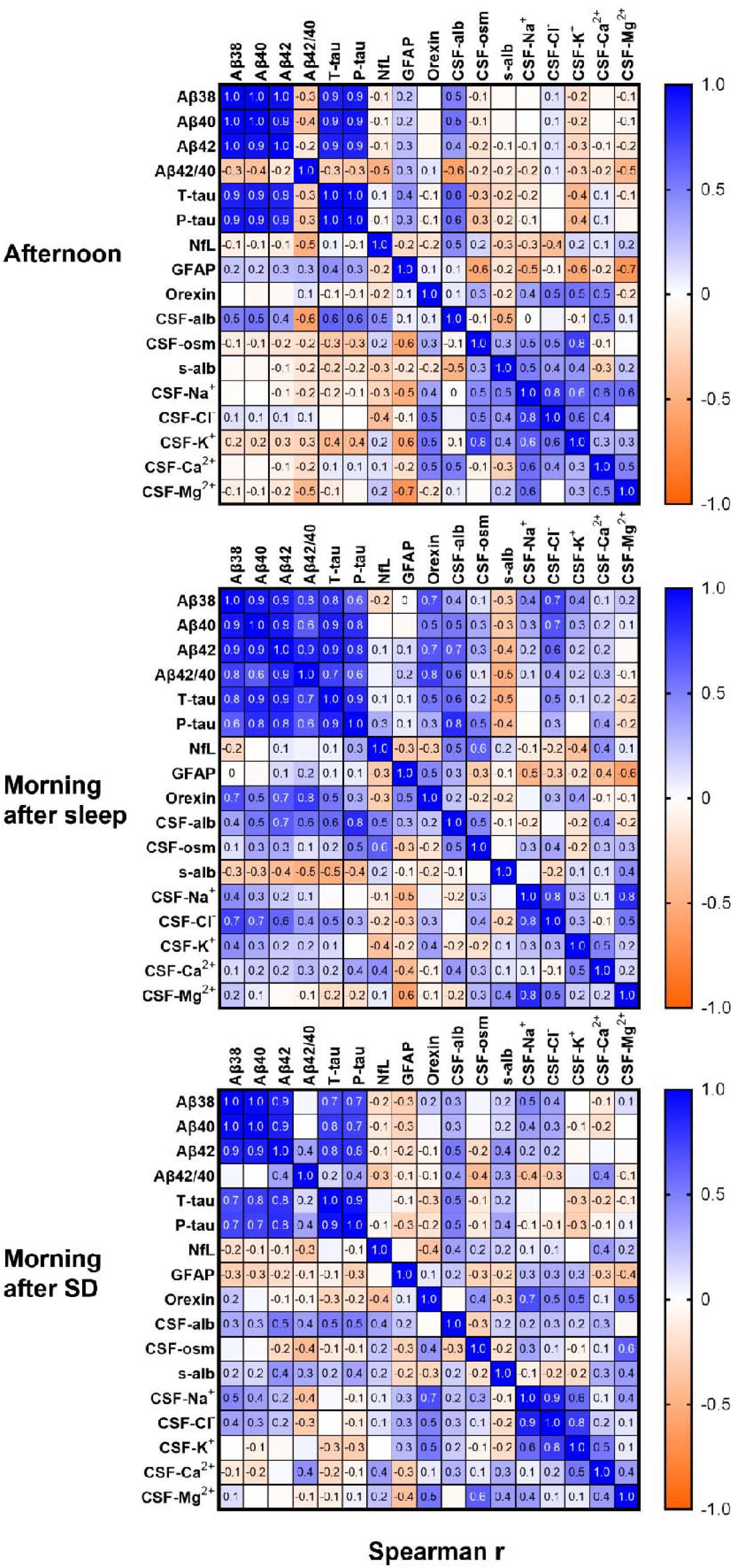
Strong correlations between Aβ and tau, but not NfL or GFAP. Spearman correlation (r) between the measured biomarkers across conditions. Abbreviations: Aβ38, amyloid β (1-38); Aβ40, amyloid β (1-40); Aβ42, amyloid β (1-42); Aβ42:40, ratio of beta amyloid 42/40; T-tau, total tau; P-tau, phosphorylated tau; NfL, neurofilament light chain; GFAP, glial fibrillary acidic protein; CSF-alb, CSF albumin; CSF-osm, CSF osmolality; s-alb, serum albumin. Data representation: Spearman correlation coefficients (r) range from -1 to 1.

### Strong correlations between Aβ and tau, but not NfL or GFAP

Patterns in the correlation analyses (Figure 3) demonstrated that CSF Aβ and tau (all the measured forms) concentrations consistently correlated strongly with each other and moderately with CSF albumin under all conditions.

Aβ and tau also showed intermediate to strong correlations with orexin, but only in mornings after sleep, while their correlations with NfL and GFAP were markedly weaker and inconsistent across conditions. CSF-ion concentrations exhibited consistent correlations with each other, separately from the Aβ and tau correlations.

## Discussion

In this randomized crossover experimental study in young and healthy volunteers, we examined how sleep affects CSF concentrations of key biomarkers reflecting AD neuropathology. We found that one night of sleep decreased CSF levels of Aβ and tau, but not NfL or GFAP, while albumin levels increased.

The selective reduction in Aβ and tau aligns with previous lumbar catheter-based studies [10, 11], in which these proteins, but not NfL or GFAP, rose significantly also during sleep and remained elevated after 24 hours. This has been interpreted as a redistribution of newly secreted proteins from the ISF to the lumbar CSF, driven by sampling-associated flow [36]. Similarly, a recent study comparing paired morning and evening CSF samples collected several weeks apart reported lower morning levels of Aβ, t-tau and 14 of 22 synaptic and endolysosomal proteins but not p-tau, NfL, or GFAP [36], suggesting analyte-specific diurnal variation.

Mechanistic evidence from rodent models indicates that both sleep [16] and synchronous oscillation of neuronal activity [22] promote Aβ clearance in rodents, whereas SWS disruption in humans increases CSF Aβ levels [9]. In our previous study of five-day partial sleep deprivation with preserved SWS, no significant changes were detected in Aβ, tau, NfL or GFAP [20], supporting the idea that SWS is specifically required for Aβ and tau clearance.

We also observed increased CSF albumin concentrations following sleep, suggesting increased CSF turnover during SWS rather than blood-brain barrier disruption, since albumin is predominantly determined by CSF flow [37]. The unchanged levels of NfL and GFAP argue against a general dilution effect and instead support a model in which increased interstitial solute mobility during sleep facilitates selective clearance. This enhanced solute mobility may facilitate receptor-mediated elimination of Aβ and tau without necessitating bulk directional flow. In contrast, the glymphatic model posits organized solute transport from the periarterial space to the perivenous space, implying more generalized and nonselective clearance.

Nonetheless, positive correlations between Aβ, tau and albumin across all conditions suggest that intraindividual variation (e.g., lower Aβ and tau, higher albumin after sleep) may contribute less to absolute biomarker concentrations than baseline interindividual differences.

Finally, orexin levels were elevated during sleep deprivation, which is consistent with our previous findings following partial sleep deprivation [20]. However, orexin showed no clear circadian rhythm and was correlated with Aβ and tau only in morning samples after sleep, suggesting that it may reflect sleep pressure rather than directly mediate acute global Aβ or tau accumulation during sustained wakefulness.

### Limitations

The modest sample size may explain why some group differences in Aβ and tau did not reach statistical significance and may also have limited our ability to detect differences in osmolality. However, overpowering an exploratory study such as ours would raise ethical concerns, given the limited incremental value of surpassing statistical thresholds relative to participant burden and resource use. Instead, we advocate interpreting our findings in context to generate novel, testable hypotheses.

The strengths of this study include its use of healthy volunteers not taking medications or drugs that affect sleep, its rigorous control of actual sleep conditions, and the theoretically low bias introduced by the sampling procedure.

## Conclusions

We propose that increased CSF turnover during slow-wave sleep selectively reduces Aβ and tau levels. Elucidating the mechanisms underlying this process may reveal novel targets for preventing or slowing pathological protein aggregation in neurodegenerative diseases.

## List of abbreviations

CSF: cerebrospinal fluid
GFAP: glial fibrillary acidic protein
AD: Alzheimer’s disease
PD: Parkinson’s disease
Aβ: beta-amyloid
TSD: total sleep deprivation
SWS: slow-wave sleep
p-tau: phosphorylated tau
t-tau: total-tau
NfL: neurofilament light chain
PSG: polysomnography
RIA: radioimmunoassay
ISEs: ion-selective electrodes
IQR: interquartile range
Aβ38: amyloid β (1-38)
Aβ40: amyloid β (1-40)
Aβ42: amyloid β (1-42)
Aβ42:40: ratio of beta amyloid 42/40

## Declarations

### Ethics approval and consent to participate

This study was approved by the Swedish Ethical Review Authority in Gothenburg (Regionala etikprövningsnämnden i Göteborg), protocol number #492-18. Each participant gave oral and written informed consent to participate in the study before inclusion. The study was conducted in accordance with the Declaration of Helsinki.

### Consent for publication

Not applicable.

### Availability of data and materials

The datasets used and/or analysed during the current study are available from the corresponding author upon reasonable request.

### Competing interests

**HZ** has served at scientific advisory boards and/or as a consultant for AbbVie, Acumen, Alector, Alzinova, ALZPath, Amylyx, Annexon, Apellis, Artery Therapeutics, AZTherapies, Cognito Therapeutics, CogRx, Denali, Eisai, LabCorp, Merry Life, Nervgen, Novo Nordisk, Optoceutics, Passage Bio, Pinteon Therapeutics, Prothena, Red Abbey Labs, reMYND, Roche, Samumed, Siemens Healthineers, Triplet Therapeutics, and Wave; has given lectures in symposia sponsored by Alzecure, Biogen, Cellectricon, Fujirebio, Lilly, Novo Nordisk, and Roche; and is a cofounder of Brain Biomarker Solutions in Gothenburg AB (BBS), which is a part of the GU Ventures Incubator Program (outside submitted work).

### Funding

The authors would like to acknowledge grants from the Swedish Research Council (VR 00986 to **EH** and #2023-00356, #2022-01018 and #2019-02397 to **HZ**), Hjärnfonden (FO2021-0048 to **EH**), Swedish State Support for Clinical Research (ALFGBG 427611 to **EH**, ALFGBG 166432 to **JH** and ALFGBG-71320 to **HZ**), Alzheimerfonden (AF-640391 to **EH**), Swedish Heart and Lung Foundation (20180585 to **JH**), Wilhelm and Martina Lundgrens Vetenskapsfond (#2018-2616 to **MF**), Göteborgs Läkaresällskap (#GLS-878091 to **MF**) and the European Union’s Horizon Europe research and innovation programme under grant agreement No 101053962 (to **HZ**).

## Authors’ contributions

Conceptualization, **MO and MF**; Methodology, **MO and MF**; Investigation, **MO, MF, HZ**, and **PW**; Formal Analysis, **TL, MO, MF**; Writing – Original Draft, **TL**; Writing – Review & Editing, **TL** and **MO**; Funding Acquisition, **HZ, JH** and **EH**; Resources, **JH**; Supervision, **HZ, PW, JH** and **EH**. All authors contributed to the interpretation of the data and critically reviewed the manuscript for important intellectual content.

## Additional files

## Supplementary Table 1

Supplementary Table 1. Medians and interquartile ranges from Figure 1 Biomarker concentrations across conditions: median and interquartile range

