## Supplementary Table 1 for "Slow Wave Sleep Reduces CSF Concentrations of Beta-amyloid and Tau: A Randomized Crossover Study in Healthy Adults"

**Supplementary Table 1. Median and interquartile ranges from Figure 1**

| | Sleep-PM | | | Sleep-AM | | | TSD-AM | | |
|---|---|---|---|---|---|---|---|---|---|
| | Median | IQR | | Median | IQR | | Median | IQR | |
| **Aβ38 (pg/mL)** | 2675 | 2507 - | 3096 | 2543 | 1933 - | 2873 | 2619 | 2393 - | 3183 |
| **Aβ40 (pg/mL)** | 7005 | 6550 - | 7923 | 7039 | 5720 - | 7596 | 7158 | 6575 - | 8251 |
| **Aβ42 (pg/mL)** | 712 | 666 - | 809 | 736 | 563 - | 799 | 793 | 648 - | 819 |
| **Aβ42/40** | 0.102 | 0.099 - | 0.103 | 0.102 | 0.098 - | 0.110 | 0.105 | 0.097 - | 0.107 |
| **P-tau (pg/mL)** | 43 | 35 - | 51 | 39 | 30 - | 41 | 44 | 33 - | 49 |
| **T-tau (pg/mL)** | 226 | 196 - | 289 | 211 | 158 - | 245 | 238 | 177 - | 261 |
| **NfL (pg/mL)** | 190 | 153 - | 253 | 200 | 180 - | 308 | 185 | 153 - | 283 |
| **GFAP (pg/mL)** | 371 | 261 - | 477 | 324 | 256 - | 440 | 414 | 286 - | 479 |
| **Orexin (pg/mL)** | 688 | 620 - | 766 | 705 | 605 - | 797 | 827 | 765 - | 939 |
| **CSF-alb (mg/L)** | 156 | 121 - | 188 | 178 | 152 - | 211 | 157 | 106 - | 179 |
| **CSF-osm (mOsm/L)** | 322 | 317 - | 329 | 318 | 312 - | 323 | 320 | 314 - | 330 |
| **s-alb(g/L)** | 43 | 41 - | 44 | 44 | 43 - | 46 | 44 | 41 - | 47 |

Biomarker concentrations across conditions.

Abbreviations: Sleep-PM, afternoon samples after nighttime sleep; Sleep-AM, morning samples after sleep; TSD-AM, morning samples after total sleep deprivation; IQR, interquartile range; Aβ38, amyloid β (1-38); Aβ40, amyloid β (1-40); Aβ42, amyloid β (1-42); Aβ42:40, ratio of beta amyloid 42/40; T-tau, total tau; P-tau, phosphorylated tau; NfL, neurofilament light chain; GFAP, glial fibrillary acidic protein; CSF-osm, CSF osmolality; s-alb, serum albumin concentration.
